# Auditory perceptual expertise: Amplitude modulation rate discrimination near the threshold for detection

**DOI:** 10.64898/2026.05.06.723339

**Authors:** Tonatiuh Garcia Ruiz, Dan H. Sanes

## Abstract

Many perceptual skills improve with a few days of training. However, weeks or months of practice may be required to reach a level of expertise on complex tasks (Watson, 1980). Here, we explored how gerbils attain expertise on a difficult task: amplitude modulation (AM) rate discrimination at very shallow AM depths, similar to the depths used during vocal communication. Using an appetitive Go-Nogo procedure, we first trained 6 gerbils to perform an AM discrimination task (Nogo: 4 Hz; Go: 4.25-10 Hz) at a depth of 0 dB (re: 100% depth). Animals were then trained to perform AM discrimination at successively shallower depths, from -3 to -18 dB, requiring an average of 5-10 days of practice to reach a performance metric of d’≥1 for each depth. Finally, we determined that AM discrimination thresholds were nearly identical between 0 to -12 dB, and only slightly elevated at -15 dB. Improvements in performance were accompanied by a large reduction in response time during procedural learning, and a gradual reduction of response time during perceptual learning, even as AM depth became shallower (i.e., more difficult). The shallowest depth at which gerbils displayed peak performance on the AM discrimination task is similar to their lowest AM depth detection thresholds. These results suggest performance on challenging auditory perceptual tasks require prolonged practice, and is accompanied by increased automaticity (i.e., lower response time) that stabilizes once expertise is achieved.

## Introduction

Acquiring new skills requires practice that refines motor, cognitive and perceptual abilities, eventually leading to expertise, defined operationally as a plateau state where consistent peak performance allows more accurate and quicker responses to the environment (Bieberman & Shiffrar, 1987; Seitz, 2020; Sheridan & Reingold, 2014; Waite et al., 2020). Expertise is fundamental to the performance of essential skills, such as prey hunting (Anholt et al., 2020). Although expertise is well-characterized for many human domains (Croijmans et al., 2020, 2021; Croijmans & Majid, 2016; Del Villar et al., 2007; Jarodzka et al., 2010; Kasarskis et al., 2001; Reingold et al., 2001; Schempp & Woorons, 2018; Sheridan & Reingold, 2014), these reports typically focus on cross-sectional data (e.g., amateurs versus experts) long after extensive practice has occurred. Alternatively, improvement on a single sensory or motor parameter is monitored over several days of practice during which performance appears to asymptote. Here, we explored how gerbils attained expertise on a difficult task that involved two fundamental parameters of amplitude modulation (AM) - rate and depth - both of which are critical for aural communication.

While expertise is found for many abilities, experts share certain characteristics. They generally perform tasks more accurately and more rapidly than novices (Bieberman & Shiffrar, 1987; Schriver et al., 2008; Waite et al., 2020). This is due to the emergence of automaticity, whereby relevant stimuli are detected efficiently, thereby reducing attentional load and decreasing response time (Newell & Rosenbloom, 1981; Reingold et al., 2001; Schneider & Shiffrin, 1977; Shiffrin & Schneider, 1977; Treisman et al., 1992). For instance, chess experts have a larger visual span than novices, permitting them to process configuration patterns that are not apparent to novices (Gauthier et al., 2003; Gauthier & Tarr, 2002). Furthermore, experts are more attentive to relevant features of the stimulus, and better able to use partial information (Hagen et al., 2023; Jarodzka et al., 2010; Sheridan & Reingold, 2014).

In the auditory domain, training-induced improvements in adult perception (i.e., perceptual learning) are thought to contribute to expertise, and have largely been described for individual acoustic dimensions (Bao et al., 2004; Beitel et al., 2003; Caras & Sanes, 2017; Kacelnik et al., 2006; Mossbridge et al., 2006; Polley et al., 2006; Recanzone et al., 1993; Van Wassenhove & Nagarajan, 2007; Wright et al., 1997; Wright & Fitzgerald, 2001). Depending on species and task difficulty, improvement is typically characterized during 1-14 daily practice sessions. Therefore, it is often uncertain whether further improvement would occur during a significantly longer period of practice. In fact, only a few experiments have measured learning during prolonged practice lasting months (Watson, 1980). Taken together, these studies suggest that training and testing on auditory psychometric tasks is generally not designed to assess the emergence of expertise.

AM detection and discrimination play a significant role in aural communication (Drullman et al., 1994; Krause & Braida, 2004). In fact, vocalization processing, including human speech, often occurs at shallow modulation depths, especially in the presence of noise (Elhilali et al., 2003; Varnet et al., 2017). Despite the importance of shallow modulation depths, with few exceptions, AM rate discrimination has only been assessed at a depth of 100% modulation (i.e., 0 dB, where depth in dB = 20*log_10_(stimulus % depth/100% depth). However, trained musicians can perform a musical interval recognition task with AM noise stimuli at close to the thresholds for AM detection itself, suggesting that long-term experience plays an important role (Burns & Viemeister, 1976; Viemeister, 1979). Therefore, we asked whether a prolonged period of training with successively more difficult conditions would permit gerbils to attain a level of expertise on an auditory task that is relevant to aural communication, AM detection at shallow depths. Our results suggest that highly trained gerbils display peak AM discrimination thresholds and rapid response times at depths near the limit of AM detection itself.

## Methods

### Experimental subjects

A total of 6 adult (3 males, 3 females) Mongolian gerbils (*Meriones unguiculatus*) were used in the study. Animals were postnatal day (P) 80 at the start of the experiment. Animals were raised from commercially obtained breeding pairs (Charles River Laboratories) and housed on a 12h light/ 12h dark cycle with full access to food and water. All procedures were approved by the Institutional Animal Care and Use Committee at New York University.

### Auditory testing environment

The animals were tested on an amplitude modulation (AM) rate discrimination task using a Go– Nogo appetitive conditioning procedure. The testing environment was a cage (dimensions: 0.4 x 0.4 x 0.4 m) that was housed in a sound attenuation booth (Industrial Acoustics; internal dimensions: 2.2 x 2 x 2 m). Stimuli were delivered from a calibrated free-field tweeter (DX25TG0504; Vifa) that was positioned 1 m above the test cage. Sound calibration measurements were made with a 1/4 inch free-field condenser recording microphone (Bruel & Kjaer). A pellet dispenser (Med Associates Inc, 20 mg) was connected to a food tray placed within the test cage, and a platform was placed on the opposite site. Both the food tray and the platform were equipped with IR emitters and sensors (Digi-Key Electronics; Emitter: 940 nm, 1.2 V, 50 mA; Sensor: Photodiode 935 nm 5 ns). Stimuli, food delivery, and behavioral data acquisition were controlled through custom MATLAB scripts and an RQ6 multifunction processor (Tucker-Davis Technologies).

### Auditory Stimuli

The auditory stimuli were amplitude modulated (AM) broadband noise tokens (4-12 Hz; 56 dB SPL; 1 s duration). All stimuli were preceded by a 200 ms onset ramp followed by an unmodulated period of 200 ms. AM depths also varied between 0 to -18 dB, re: 100% depth (where depth in dB = 20*log_10_ (stimulus % depth/100% depth) in 3 dB intervals.

### Procedural learning

Gerbils were placed on controlled food access and trained on the procedure using a social learning paradigm that ran for 14 days (Paraouty et al., 2020, 2021, 2023). Animals were initially paired for 5 successive days with a trained demonstrator that performed the AM discrimination task. Following this social exposure, gerbils were placed alone in the test cage and permitted to practice the AM discrimination task for 9 days. This required that the gerbil learn to initiate a trial by standing on a platform to interrupt an infrared beam, and approach a food tray upon presentation of the Go stimuli (AM rates of 6, 8, or 10 Hz) where they received a reward (two 20 mg pellets per trial; BioServ) delivered from a pellet dispenser (Med Associates Inc.). They also learned to withhold a response to approach the food tray upon presentation of the Nogo stimulus (4 Hz AM).

Trials were scored as a Hit (correctly approaching the food tray during a Go trial), Miss (failing to approach the food tray), Correct Reject (CR; correctly reinitiating a trial during a Nogo trial), or False Alarm (FA; incorrectly approaching the food tray on a Nogo trial). For false alarms, no food was delivered and the house lights were turned off for 8 sec during which a trial could not be initiated. Hit and FA rates were constrained to floor (0.05) and ceiling (0.95) values to avoid d′ values that approach infinity. Psychometric threshold was defined as the AM rate at which d′ = 1. Performance was quantified using the signal detection metric, d’, defined as d’ = z(hit rate) – z(false alarm rate).

### Perceptual learning of AM rate discrimination successively shallower AM depths

During perceptual learning, animals were typically presented with 3 AM rates on Go trials (5, 6, and 8 Hz) and the 4 Hz Nogo. Since animals had already been tested at 0 dB during procedural learning (above), they were first trained with an AM depth of -3 dB. Each animal’s FA rate, number of trials per session, and d’ were monitored across depths, and these values were used to determine whether an animal could advance to the next shallower depth. The general goal was for each depth was to attain a d’≥1, a FA rate <40%, and ≥100 trials per session. Once animals achieved a d’≥1, AM depth was reduced by 3dB. This training process was repeated for depths of -6, -9, -12, -15 and -18 dB. At each depth, animals were trained with 2 procedures: (1) During interleaved sessions, animals were trained with 2 contiguous depths (e.g. 0dB/-3dB, -3dB/-6dB). (2) During blocked sessions, animals were trained on a single depth (Supplemental Figure 1). The perceptual training ran for 39-50 days, depending on each gerbil’s performance (Supplemental Figure 1).

**Figure 1.**
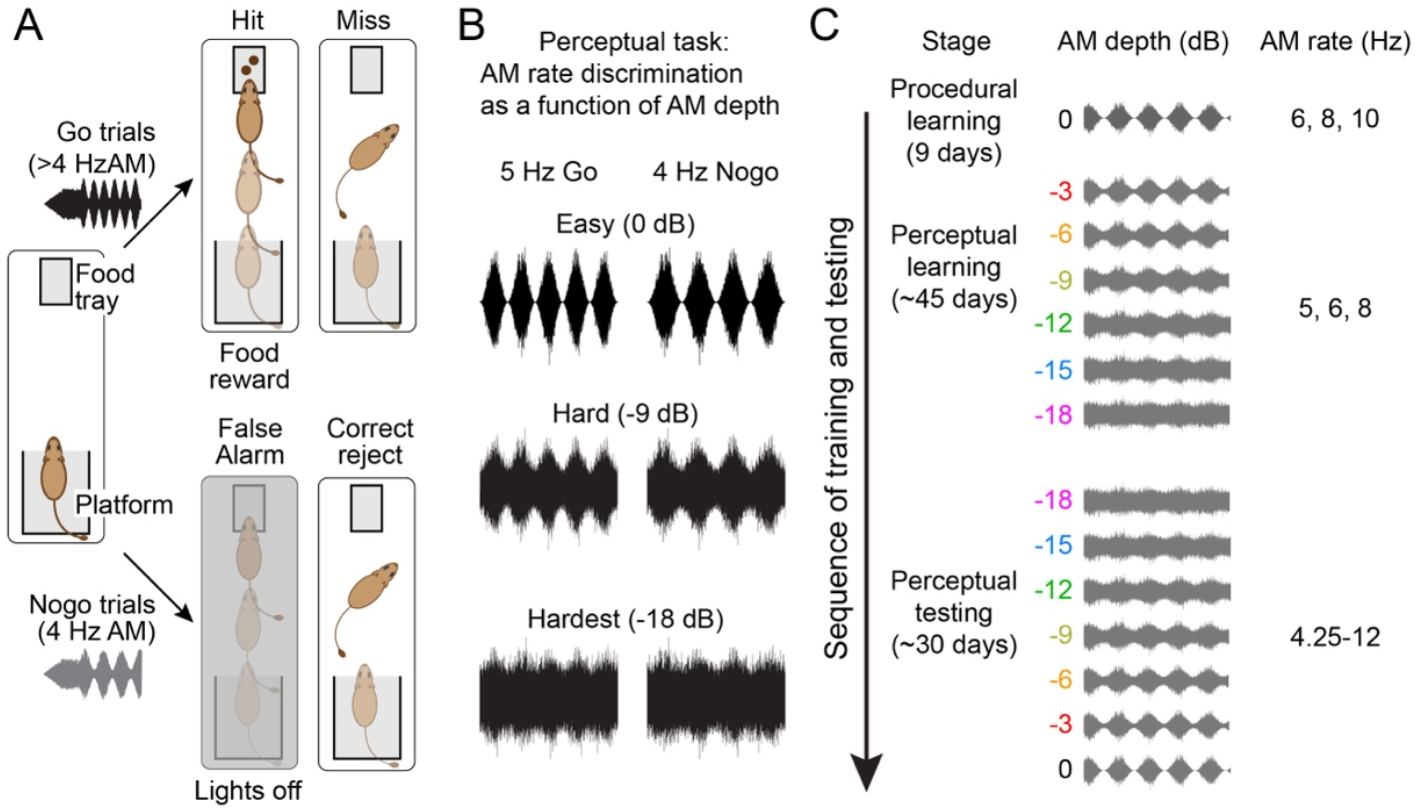
Methods. Animals were trained and tested in an AM discrimination task at low frequencies and decreasing depth. **(A)** Go-Nogo task. When the animal started a trial by standing on the platform, it had 5 s to reach the food tray. If the stimulus is a >4Hz sound and gets to the food tray within time, it is computed as a Hit and food is given. If the animal doesn’t reach the food tray, the trial is computed as a miss, and the animal doesn’t receive food. On the other hand, if the stimulus is a 4hz sound and it reaches the food tray within 5 seconds, the trial is computed as a False alarm (FA) and the animal is punished with a time out of 8 seconds with the light off. Similarly, if the animal abstains from reaching the food tray within time, it is computed as a correct reject (CR). **(B)** Stimuli waveform examples of 5hz and 4hz stimuli at 0, -9 and -18dB. **(C)** Sequence of training for the experiment, including AM depths and AM rates presented at each stage.

### Perceptual testing of AM discrimination thresholds at each AM depth

Once animals performed the AM discrimination task at the full range of depths, 0 to -18 dB, we obtained their AM rate discrimination thresholds at each depth. Animals were first tested at -18dB and then at 3 dB increments using both interleaved and blocked sessions until we obtained a threshold per depth. Therefore, we introduced lower AM rates, down to a minimum of 4.25 Hz, when necessary. The perceptual testing ran for 25-33 days, depending on each gerbil’s performance.

## Results

We explored the acquisition and expert performance in AM discrimination at shallow depths in 3 phases: (1) Procedural training during which animals learned to perform the AM discrimination task at a depth of 0 dB (Figure 2), (2) perceptual training during which animals learned to perform the AM discrimination task at successively shallower AM depths (Figure 3), and (3) perceptual testing during which we obtained AM discrimination thresholds at each modulation depth (Figure 4). The total number of sessions ranged from 83-90 days, and the total number of trials ranged from 8,500-12,000.

**Figure 2.**
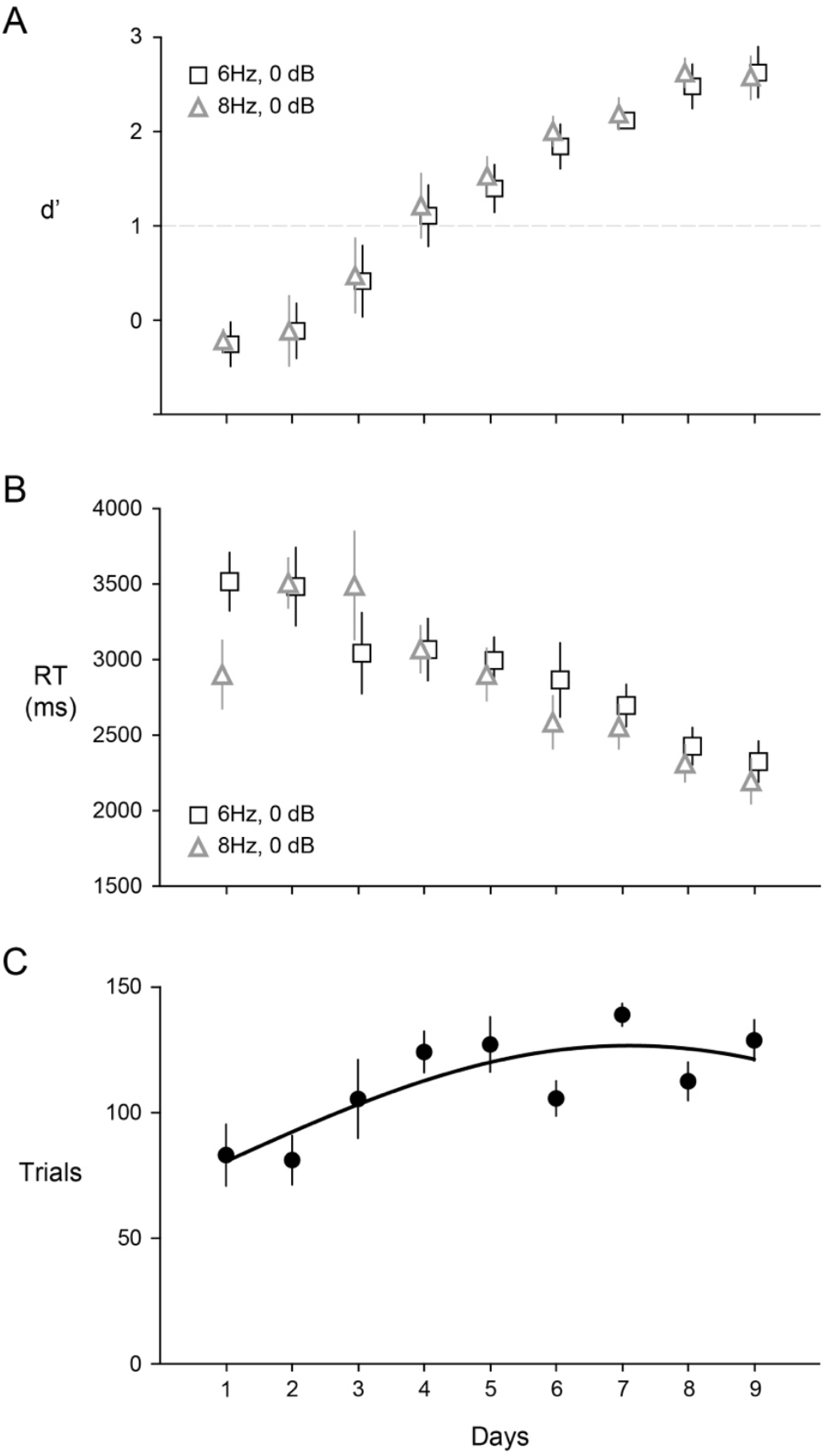
Procedural learning. Animals first learn to perform the task using a social learning paradigm, where after social exposure, animals are left to practice the task for 9 days at 0dB. **(A)** 6Hz, 8Hz d’ show an exponential improvement in performance within 9 days. Average d’ (± SEM) is reached at day 4 of practice and it plateaus over d’=2 by day 9. **(B)** Response time (± SEM) steadily decreased more than 1000 ms within the 9 days of practice. **(C)** As days of practice advance, animals increase the number of trials/session (± SEM) by an average of 45.6 from beginning to end of the procedural learning.

**Figure 3.**
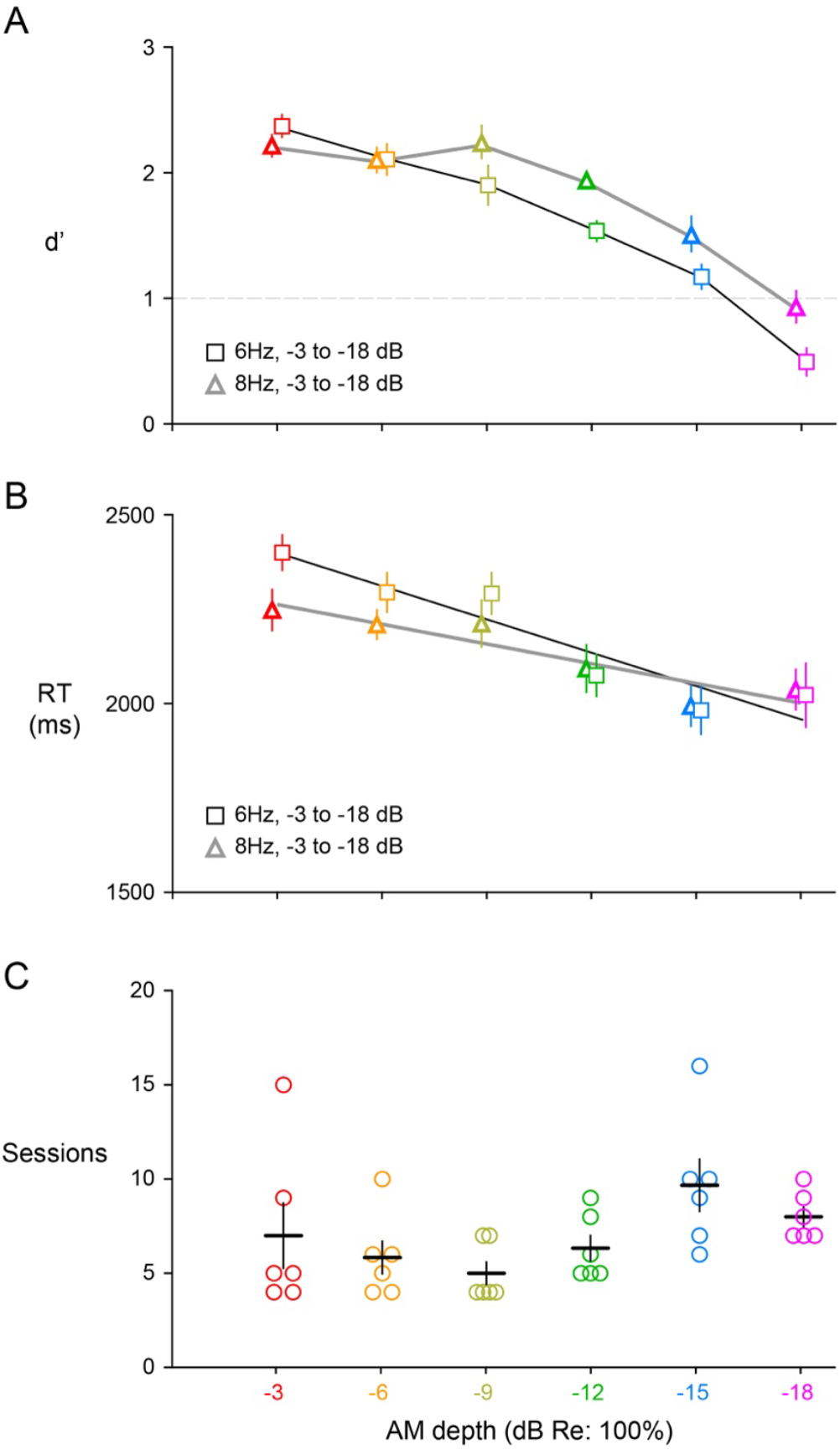
Perceptual learning. (**A**) Average discriminability performance (d’) for 6 and 8hz (± SEM) during perceptual learning for sessions for best sessions. Animals decrease performance as depth becomes shallower, having -18dB reach a d’<1; this shows how hard the task becomes for the animal (**B**) Response time (± SEM) continues decreasing by more than 300ms even when the task becomes harder and harder. (**C**) Number of training sessions necessary to reach criterion per depth (± SEM). A higher number of sessions are necessary at shallower depths.

**Figure 4.**
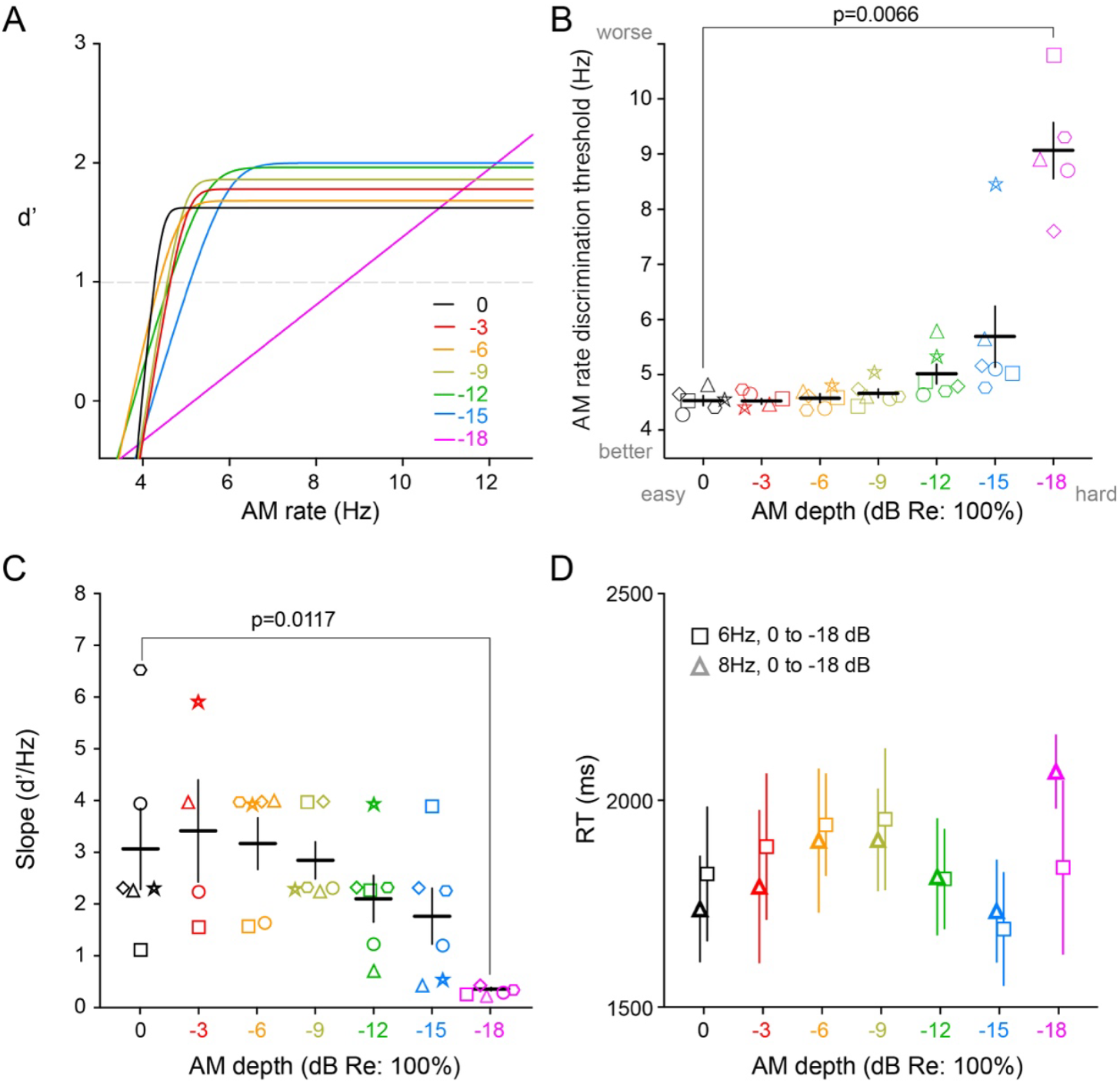
Perceptual testing. (**A**) Psychometric curve for one animal at different depths. Animal threshold is defined where the line crosses d’=1. (**B**) AM rate discrimination thresholds per depth. From 0dB to -15dB, the threshold is very similar but it dramatically increases at -18dB. One-way ANOVA with Dunnett test indicates significant differences only between 0dB and -18dB (F(2.037, 9.167)=29.92, p<0.0066). 0dB (4.5 ± 0.08), -3dB (4.52 ± 0.06), -6dB (4.57 ± 0.07), -9dB (4.66 ± 0.081), -12dB (5.01 ± 0.18), -15dB (5.69 ± 0.56), -18dB (9.06 ± 0.52) (± SEM) (**C**) Psychometric curve slope per depth. Significantly shallower slopes are found at -18dB. 0dB (3.07 ± 0.78), -3dB (3.41 ± 0.97), -6dB (3.17 ± 0.50), -9dB (2.84 ± 0.36), -12dB (2.12 ± 0.45), -15dB (1.76 ± 0.54), - 18dB (0.30 ± 0.033) (± SEM). Linear mixed-effects model (REML), followed by a Holm-Šidák multiple comparison test show no significant differences between 0dB and -18dB (F(1.973, 8.878)=3.506, p=0.0759). (± SEM). One-way ANOVA with Dunnett test show only significant differences between 0dB and -18dB, showing no significant differences in slope at the other depths. **(D)** Response time stabilizes and varies across depths (± SEM).

### Procedural learning to acquire the AM discrimination task

Following social exposure to a trained demonstrator (see Methods), animals practiced the AM discrimination task at a depth of 0 dB for 9 days. Animals reached a d’=1 by day 4 (n=6), and reached peak performance of d’ of 2.5 at day 8 (Figure 2A). As animals improved on the task, their response time (RT) decreased significantly by an average of 1192 ms (Figure 2B). The number of trials also increased as animals learned the task and got better performance by an average of 45.6 trials from beginning to end (Figure 2C).

### Perceptual learning of AM rate discrimination on shallow AM depths

Once animals completed procedural learning on the AM discrimination task at 0 dB with multiple Go stimuli (6, 8, 10 Hz), we asked whether they could generalize to different AM rates and shallower AM depths. When tested at a depth of 0 dB, animals were immediately able to perform the task with each of the lower AM rates: 5, 6 and 8 Hz. However, animals required additional practice to perform the AM task after introducing them to each of the shallower AM depths, and each animal displayed a unique learning trajectory (Supplemental Figure 1). We observed that d’ decreased for all tested AM rates as animals were trained on successively shallower AM depths (Figure 3A). This effect was more apparent at lower rates and shallower depths. For the most challenging AM rates, FA rates increased and hit rates decreased (Supplemental Figure 2). During perceptual learning, RT continued to decline across practice sessions, even though the task became increasingly difficult at shallower AM depths (Figure 3B). This suggests that the sensory decision and motor output continue to improve which is consistent with an improvement in automaticity. Figure 3C shows the number of sessions required before advancing to the next shallower depth (-3 to -18 dB). Animals required an average of 4-10 sessions to reach criterion at each depth. (Figure 3C). The entire training period for perceptual learning lasted an average of 43 ± 4 days (Supplemental Figure 1). Finally, criterion values tended to become more negative (e.i., animals displayed a more liberal response) during the course of perceptual learning and perceptual testing, as the task became more difficult (Supplemental Figure 2).

### Perceptual testing of AM discrimination thresholds at each AM depth

Once animals completed perceptual learning, we obtained their AM discrimination thresholds at each AM depth. Figure 4A shows representative psychometric functions from one animal, and illustrates that similar performance was observed from 0 to -15 dB. Figure 4B shows each individual animal’s AM discrimination threshold at d’=1. There was no significant difference between discrimination thresholds from 0 to -15 dB, but thresholds increased significantly at -18 dB (Linear mixed-effects model (REML), followed by a Holm-Šidák multiple comparison test: F(2.037, 9.167)=29.92, p<0.0066). We note that one poor performing gerbil largely accounted for the decline in threshold at -15 dB, and this animal was unable to perform the task at -18 dB. Psychometric function slope did not differ significantly between 0 to -15 dB, but was shallower at -18dB. This suggests that animals were less confident at -18dB. Finally, Figure 4D shows that there were no significant differences in RT across all depths tested (Linear mixed-effects model (REML), followed by a Tukey multiple comparison test: F(1.647, 7.961)=1.152, p=0.3512). This suggests that automaticity stabilized when animals reached a level of expertise.

## Discussion

Rapid and accurate processing of the subtle acoustic features that compose speech, including AM discrimination, are critical for aural communication (Drullman et al., 1994; Krause & Braida, 2004). This set of skills requires years of experience, suggesting prolonged learning and the emergence of expertise. For example, the categorization of voice emotion or recognition of talker identity continue to improve through early adolescence (Amorim et al., 2021; Grosbras et al., 2018; Levi, 2018; Levi & Schwartz, 2013; Mann et al., 1979; Rachman et al., 2025). In contrast, the emergence of peak performance is typically documented at a single time point through cross-sectional comparisons between experts and amateurs (Gallo et al., 2025; Mischler et al., 2025; Rigoulot et al., 2015; Varnet et al., 2015). To address the emergence of expertise, we characterized the acquisition of an AM rate discrimination task at shallow modulation depths, a skill that is relevant to speech processing, especially in the presence of noise (Elhilali et al., 2003; Liu & Zeng, 2006; Varnet et al., 2017). With prolonged practice, gerbils displayed peak rate discrimination thresholds between depths of 0 to -12 dB, and only slightly elevated thresholds at -15 dB which is the approximate limit for AM detection, itself (Anbuhl et al., 2022; Caras & Sanes, 2015; Masri et al., 2024; Rosen et al., 2012; Von Trapp et al., 2016). Furthermore, the acquisition of peak perceptual performance was accompanied by a reduction in response time (RT), a hallmark of expertise (Newell & Rosenbloom, 1981; Reingold et al., 2001).

We explored the acquisition and expert performance in AM discrimination at shallow depths in 3 phases: Procedural learning, perceptual learning, and perceptual testing. Depending on the animal, the entire training period ranged from 83-90 days, and used 8,500-12,000 trials. Procedural learning led to baseline proficiency on the AM discrimination task. AM rate discrimination performance displayed a steep learning curve during 9 days of practice (Figure 2A), and this was accompanied with a dramatic decrease of RT of more than 1000 ms (Figure 2B). Steep learning curves have been described as a period of effortful learning related to procedural memory consolidation (Hong et al., 2019). Thus, procedural learning provides animals with general task proficiency, including a significant decrease in RT that is associated with a process of automaticity (Newell & Rosenbloom, 1981; Reingold et al., 2001).

During perceptual learning, gerbils practiced the AM discrimination task at successively shallower depths between 0 to -18 dB. Peak performance, operationalized as the d’ value observed for two AM rates (6 and 8 Hz) became poorer as the AM depth became shallower. However, gerbils learned to perform the task through a depth of -15 dB (Figure 3A). Despite increasing task difficulty, as AM depth became shallower, the RT continued to decline by approximately 400 ms (Figure 3B). In fact, experts demonstrated heightened sensitivity and faster access to task-relevant features which may contribute to shorter response times (Jarodzka et al., 2010; Sheridan & Reingold, 2014). Therefore, the reduction in RT observed here suggests a further extension of automaticity (Newell & Rosenbloom, 1981; Reingold et al., 2001). In principle, as gerbils attain expertise on the AM discrimination task, they are better able to attend to subtle, but important, features of the AM envelope. In fact, eye tracking studies have shown that quicker and more accurate responses from experts is accompanied by changes in attention that allow subjects to identify important features in the task in hand (Bieberman & Shiffrar, 1987; Schriver et al., 2008; Sheridan & Reingold, 2014; Waite et al., 2020).

During perceptual testing, gerbils ultimately displayed AM rate discrimination thresholds of 4.6 ± 0.2 Hz at -12 dB, nearly identical to AM discrimination thresholds at 0 dB depth (current study Figure 4B: 4.5 ± 0.1; (Von Trapp et al., 2017): 4.8 ± 0.3 Hz; (Yao & Sanes, 2021): 4.9 ± 0.2 Hz). Furthermore, AM discrimination was only marginally poorer at -15 dB (5.7 ± 0.2 Hz), but became significantly worse at -18 dB (Figure 4B). Finally, the RT observed during perceptual testing stabilized at, or below, the lowest durations observed during perceptual learning, indicating that automaticity was sustained despite the task once again becoming more difficult in the AM rate domain (i.e., additional smaller AM rates were added to the task) (Figure 4D). The human literature suggests that AM discrimination at low rates (i.e., <70 Hz) requires large AM depth, as compared to AM detection thresholds themselves (Patterson et al., 1978; Viemeister, 1979). However, trained musicians can perform a musical interval recognition task with AM noise stimuli at close to the thresholds for AM detection (Burns & Viemeister, 1976). Therefore, our results are consistent with observations from human listeners which suggest that expertise is needed for AM discrimination at shallow depths.

It is important to address whether our results should be thought of as an instance of perceptual learning or perceptual expertise. It has been difficult to compare these two phenomena because they typically have different experimental designs. Perceptual learning studies tend to focus on learning rates for a single perceptual feature over a relatively short period of time (Caras & Sanes, 2019; Demany, 1985a; Glennon et al., 2019; Kattner et al., 2017b; Klingel, Singhal, Seitz, & Kopco, 2025; Watson, 1980). In contrast, studies of expertise examine complex tasks with a cross-sectional design that compares experts to novices (Busey & Vanderkolk, 2005; Hagen et al., 2023; McKone et al., 2006; Xu et al., 2016). We interpret our results to suggest that extensive perceptual learning leads to perceptual expertise because gerbils ultimately performed one task (AM discrimination) at depths close to the limit of AM detection, and displayed a steady decline in response time even as the task became more difficult (Figure 3), stabilizing during perceptual testing (Figure 4). Our results are consistent with human findings in which a comparison was made between a perceptual learning and perceptual expertise task using the same stimuli, revealing that individuals displayed similar characteristics on generalization, another hallmark of expertise (Wong et al., 2011).

Our experiment considered the emergence of expertise only in adult gerbils, yet experience and practice are a hallmark of the prolonged maturational improvement of perceptual skills (Sanes & Woolley, 2011). For example, the detection threshold for amplitude modulations (AM) matures in adolescence (Banai et al., 2011). In fact, learning itself continues to mature into adolescence (Decker et al., 2016; Huyck & Wright, 2011, 2013; Knoll et al., 2016; Lukács & Kemény, 2015; Pattwell et al., 2012). The relevance of perceptual development to speech processing is also suggested by the strong correlation between hearing loss-induced deficits in spectral modulation detection and phonological sensitivity (Nittrouer et al., 2025). Therefore, our paradigm provides only a partial assessment of expert performance because maximal expression commonly occurs when a tremendous amount of practice is distributed over years of development.

## Abbreviations

(AM): amplitude modulation
(dB): decibel
(CR): correct reject
(FA): false alarm
(RT): response time
(d-prime): d’

## Acknowledgements

This research was supported by R01DC020279 (DHS) and T32MH019524 (TGR).

## Figures

**Supplemental Figure 1.**
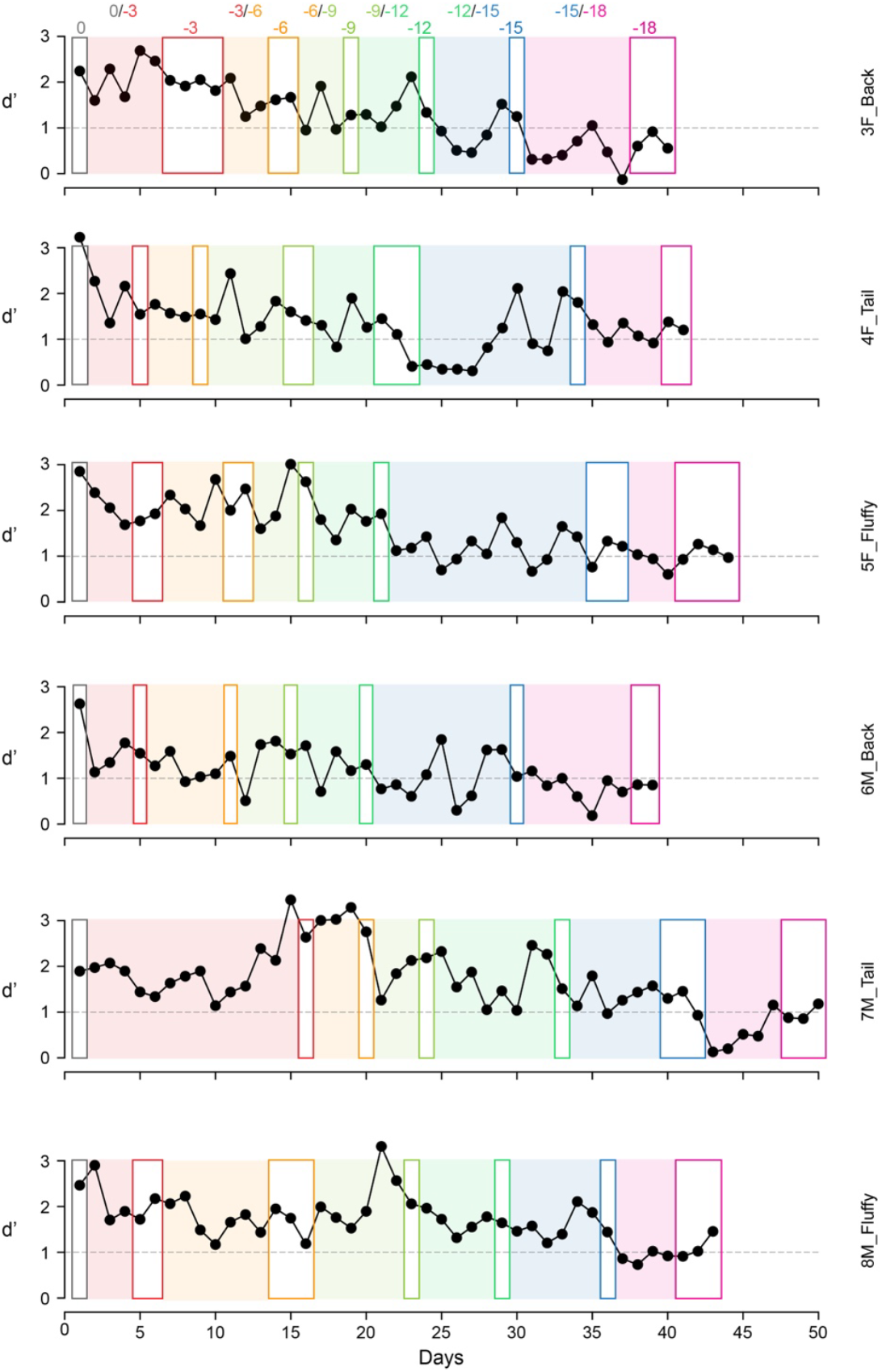
Individual animal performance during perceptual learning. Each graph presents the performance of one animal during each day of perceptual training, as we decreased the AM depth from 0 to -18 dB. Within clear boxes, animals only performed the task with a single depth, and within shaded boxes, animals performed the task with two depths. Closed symbols represent days on which animals met our performance criteria, and open symbols represent days on which animals failed to meet our criteria (see Methods).

**Supplemental Figure 2.**
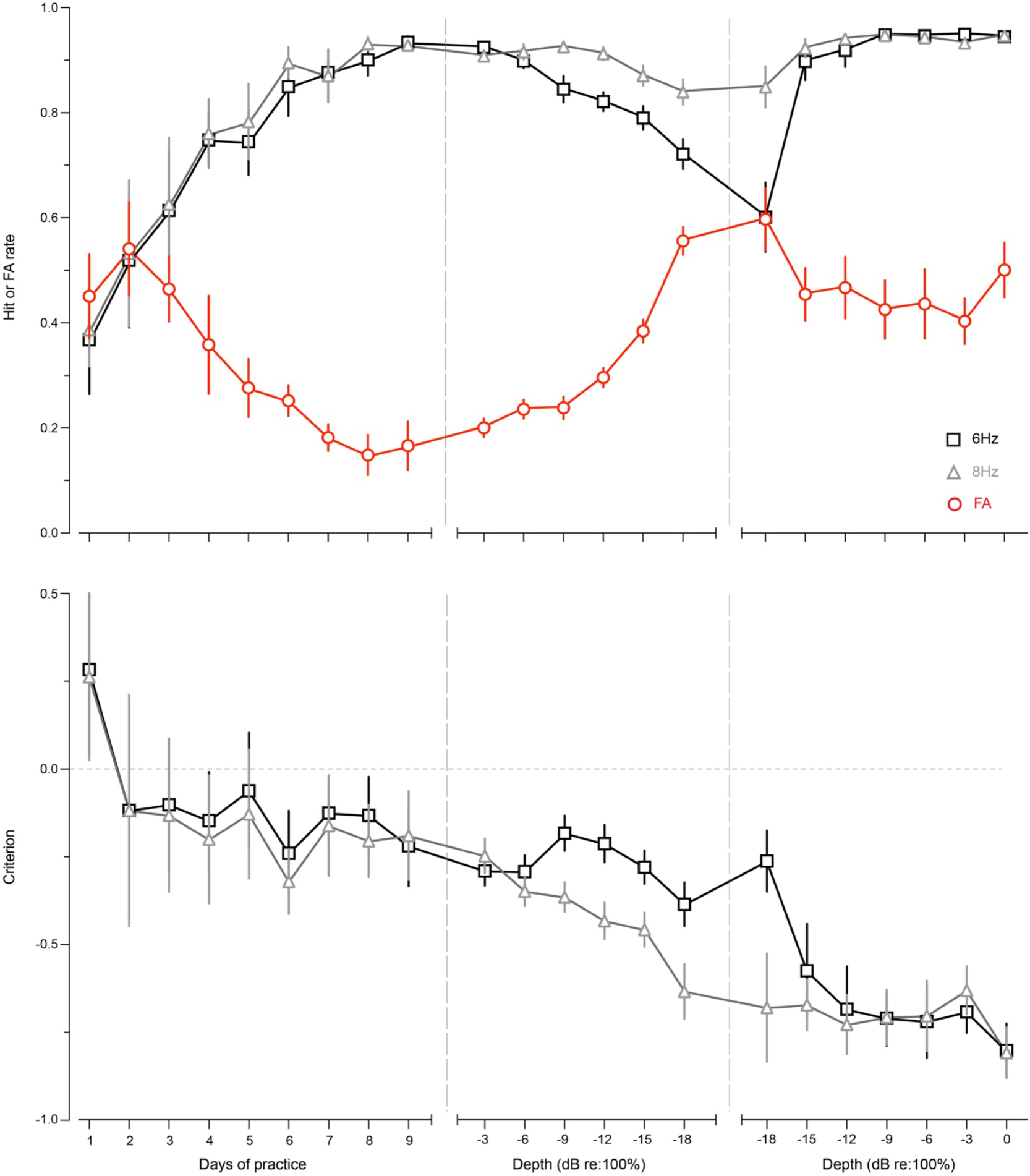
Performance metrics across the entire training period. **(A)** Hit and FA rate changes for each phase of the training and testing regimen. (**B**) Signal detection theory criterion for 6hz and 8hz for each phase of the training and testing regimen.

